# Understanding changes in genetic literacy over time and in genetic research participants

**DOI:** 10.1101/2022.09.12.507568

**Authors:** India D. Little, Laura M. Koehly, Chris Gunter

## Abstract

As genomic and personalized medicine becomes mainstream, assessing and understanding the public’s genetic literacy is paramount. Because genetic research drives innovation in this area and involves much of the public, it is equally important to assess its impact on genetic literacy. We designed a survey to assess genetic literacy in three ways (familiarity, knowledge, and skills) and distributed it to two distinct samples: 2050 members of the general population, and 2023 individuals currently enrolled in a large-scale genetic research study. We compared these data to a similar survey implemented in 2013. The results indicate that familiarity with basic genetic terms in 2021 (M = 5.36, p<.001) and knowledge of genetic concepts in 2021 (M = 9.06, p = 0.002) is significantly higher compared to 2013 (M = 5.08; M = 8.72). Those currently enrolled in a genetic study were also significantly more familiar with genetic terms (M = 5.79, p<.001), more knowledgeable of genetic concepts (M = 10.27, p<.001), and scored higher in skills (M = 3.57, p<.001) than the general population (M = 4.98; M = 9.06; M = 2.65).The results suggest that genetic literacy is improving over time, though there is room for improvement. The data also suggest that participating in genetic research is one avenue for improving genetic literacy. We conclude that educational interventions are needed to ensure familiarity with and comprehension of basic genetic concepts. We also suggest further exploration of the impact of genetic research participation on genetic literacy to determine mechanisms for potential interventions.

## Introduction

Genomic research advancements in the last decade and the rapid reduction in genomic sequencing costs have exponentially increased the use of genetic information in everyday life. The cost of whole genome sequencing has declined from just under $1 million in 2008 to as low as $17,000 in 2021, expanding opportunities for clinical genetic research and direct-to-consumer genetic testing.^1,2^ Increasing numbers of individuals are pursuing ancestry testing, clinical genetic testing, or more personalized testing options on their own.^3^ A concurrent rise in research involving genome or exome sequencing also exposes individuals to their own genetic data and its implications.^4^ This increase in the accessibility of genetic information, both within and beyond the healthcare setting, requires the public to interpret genetic testing results and apply them to their own health. To make educated decisions based on genetic information, individuals must have a basic understanding of genetic concepts and be able to effectively communicate with their providers about testing options and results. As the use of genetic information in healthcare increases, there is a need to measure and assess how individuals learn and communicate this information. Genetic literacy is one construct that can measure this ability.

Genetic literacy is defined as the sufficient knowledge and understanding of genetic principles for individuals to make decisions that sustain personal well-being and effective participation in social decisions on genetic issues.^5^ Distinct from genetic knowledge, genetic literacy evaluates both an individual’s understanding of basic genetic concepts and their ability to apply this understanding to health decisions. Genetic literacy can improve communication between patients and providers as well as help patients make informed decisions for their health.^6,7,8^ Higher genetic literacy can also improve attitudes towards genetic tests, such as the belief that genetic testing can provide important information to family members.^9^ Genetics education can also improve understanding of the limitations of genetic testing and its predictive ability, helping individuals make informed decisions about testing options and participation in genomic research.^4^ In addition, individuals with low genetic literacy benefit the least from advancements in genomic and personalized medicine and are less likely to participate in genetic research.^10^ Improving literacy in overlooked populations is crucial to limiting gaps in clinical opportunity and quality of care.^6,11^ Low genetic literacy acts as a potential threat to effective translation of genetic tests and is also associated with higher rates of heath complications, furthering health disparities.^9,11^

Despite the increase in genetic information in healthcare, medical training in genetics is often lacking, directly impacting genetic literacy. Low genetic literacy can be detrimental to quality of care and impair providers’ ability to effectively discuss genetic contributions to health with patients. A genetics knowledge survey distributed to first year pathology residents indicated significantly reduced understanding of genetics concepts compared to non-genetics concepts. Further, only 53% of the sample reported interacting with a medical geneticist during medical school.^12^ In a sample of 10,303 physicians, only 29% reported they received education in pharmacogenetic testing;^13^ many medical students also self-report insufficient genetics knowledge.^14^ The consequences of low genetic literacy in providers include misdiagnosis, treatment failure, or unnecessary genetic testing, and represent a significant barrier to the implementation of precision health medicine.^15,16^

Similarly, the general population in the US is often not adequately trained in basic genetic science. A survey distributed to 5,404 participants in the US with secondary education indicated that only 1.2% of the sample answered all 18 basic genetic knowledge questions correctly.^10^ Similar trends are seen in educators with 25% of high school teachers reporting teaching contemporary issues in genetics.^4^ Another national survey conducted in 2017 indicated that only half of individuals are aware of genetic testing and approximately a third are aware that genetic testing can contribute to disease treatment.^17^ Over 30% of a 2,093-person national sample indicated that they were unfamiliar with genetic concepts, while only 20% indicated they had personal experience with genetic health issues.^18^ Despite generally low genetic literacy in both patient and provider populations, interventions that involve and engage the public in genomic science can help assuage the disparity.^8,45^

Participating in genetic research is one way for individuals to gain exposure to genetic contributions to health, potentially despite insufficient education. Genome-wide association studies (GWAS) are a popular tool in genetic research, used to analyze associations between quantitative traits and known genetic variants.^1^ Funding within genomic research is also increasing. Since the start of the Human Genome Project, research funding provided by the National Human Genome Research Institute (NHGRI) has increased tenfold.^19^ While the practice of involving participants in genetic research is growing, its impact on participants’ genetic literacy is not fully understood. In Japan, promotion of genetic research is associated with higher genetic literacy.^20^ In addition, the informed consent process can improve participants’ understanding the of the limitations and benefits of genome sequencing.^21^ Despite increased interest and participation in genetic research, we do not fully understand genetic literacy rates in the general population compared to those currently enrolled in genetic research.

The tools used to measure genetic literacy vary, including pronunciation of health jargon, awareness of common genetic terms, and accuracy in factual knowledge.^9,22,23^ The Rapid Estimate of Adult Literacy in Genetics (REAL-G) assesses individuals’ ability to recognize and recite genetic terms within the clinical context.^22^ Similarly, the Genetic Literacy And Comprehension (GLAC) measure presents eight terms and asks participants to rate their level of familiarity.^9^ In contrast, the Genetic Literacy Assessment Instrument (GLAI) and the UNC Genomics Knowledge Scale (UNC-GKS) assess factual genetic knowledge.^23,24^ The Genomics Knowledge Scale (GKnowM) is a recently developed tool of 26 multiple-choice questions meant to assess knowledge of recent genomic developments and content needed to make informed health decisions.^25^ While all established measures assess an individual’s knowledge and awareness of genetic concepts, there is a need for a standardized measure that can be applied to improve genetic literacy.

In 2013, a comprehensive survey assessing genetic literacy and its relationship to breast cancer risk was administered to a nationally representative sample.^5^ The survey measured genetic literacy in three facets: familiarity with genetic terms (e.g. heredity, chromosome); accuracy in genetic knowledge (e.g. Is a gene bigger than a chromosome?); and sufficient skills to synthesize information applying genetic information to human health (e.g. What is the purpose of genetic testing?). These facets are based on previously used measures, and in conjunction, evaluate an individual’s awareness and understanding of basic genetic concepts and their ability to apply this information to a clinical example. Because of genetic research advances in the last decade and the drastic increase in genetic testing, there is a need to update genetic literacy rates in the general population, with consideration of how genetic literacy may relate to common, complex conditions such as autism spectrum disorder (ASD).

Here we take advantage of elements of the Abrams et al. (2015) survey to longitudinally assess genetic literacy as well as its relationship to a different common complex genetic condition. We assess genetic literacy in the same three facets by replicating the familiarity and knowledge segments, while adapting the skills segment to address genetic susceptibility for autism spectrum disorder (ASD). We have two primary aims. The first is to update genetic literacy rates for the general population and compare results to those in Abrams et al. (2015). The second is to assess genetic literacy in a population currently enrolled in a genetic research study specifically focused on ASD, called the Simons Powering Autism Research or SPARK study.^26^ Our hypotheses are that genetic literacy is measurably higher since the first administration of the survey, and that individuals participating in a genetic research project will have higher genetic literacy.

## Material and Methods

We implemented the Genetic Literacy Survey (GLS) to assess individuals’ genetic knowledge and ability to apply genetic information to human health decisions. It was developed from the survey used in Abrams et al. (2015), including the same three assessments of genetic literacy: familiarity, knowledge, and skills. The survey was completed entirely online and designed to be completed in approximately 25 minutes. The research protocol was deemed exempt by the NIH Office of Research Protections.

### Language Choices

We note that terms used around autism are fluid and personal, and whenever possible should depend on what those involved in the research prefer when describing themselves.^27,28^ For this study, we will use “autism,” “autism spectrum disorder” or “ASD” when describing the condition and “autistic” when describing individuals.

### Participants and Survey Development

#### 2013 General Population (GP) Sample

A previous version of the GLS described in Abrams et al. (2015) was distributed to a sample (N = 1016) recruited through a third-party contractor. See Abrams et al. (2015) for further details.^5^

#### 2021 General Population (GP) Sample

The GLS was distributed to a sample (N = 2050) recruited from a respondent panel compiled by a third-party contractor. The respondent panel is a group of people who have indicated their interest in completing surveys and have been prequalified, both by indicating their willingness to participate and by providing their geographic, demographic, sociographic and psychographic data. The final sample was designed to represent the general population in age, gender, and education level. The sample over-represents Black participants (27.5%) to replicate the sample distribution in Abrams et al. (2015). No participants were excluded, resulting in a final sample of 2050. Participants in the general population sample completed the electronic survey between April 12, 2021 and April 27, 2021.

#### SPARK (Genetic Study) Sample

The GLS was distributed to a sample (N = 2264) identified through the SPARK (Simons Foundation Powering Autism Research) study, a large-scale database of autistic individuals and families with autism. Individuals in this database freely elect to share their medical, genetic, and demographic data with SPARK and voluntarily completed the GLS survey through the Research Match platform. Two hundred and forty-one participants were excluded from analysis because they did not complete the survey in its entirety, resulting in a final sample of 2023. Again, the sample over-represents Black participants (20.5%) to replicate previous sample distributions. Participants in the genetic study sample completed the electronic survey between September 20, 2021 and November 21, 2021.

### Survey Measures

#### Term familiarity

Respondents were asked to rate their familiarity with eight terms – genetic, chromosome, susceptibility, mutation, variation, abnormality, heredity, sporadic – on a scale of 1 (not at all familiar) to 7 (completely familiar). Scores were calculated as an average familiarity across the eight items, with an average Cronbach’s alpha of 0.923. A variation of this scale has been used in previous measures of genetic literacy. We replicated the Genetic Literacy and Comprehension (GLAC) measure, which presented the eight common genetic terms and asked respondents to rate their level of familiarity.^9^ The short version of the Rapid Estimate of Adult Literacy in Genetics (REAL-G) also uses the same eight terms with established face and predictive validity.^22^ We added a ninth term “genome” to assess awareness surrounding recent advances in genomic medicine, including the Human Genome Project.^29^ Familiarity with this ninth term is not included in the final average familiarity score.

#### Factual Knowledge

Respondents were presented with sixteen technical genetic statements and asked to respond with “True,” “False,” or “Don’t Know” for each statement, seven of which were intentionally false. The statements concerned genes, their function and makeup, and potential risk for genetic disease. This measure replicated that used in Abrams et al. (2015).^23,30,31^ Participants received a score of 1 for correct responses and a score of 0 for incorrect or unsure responses, creating a final score range of 0 to 16. Based on Kuder-Richardson 20, the assessment demonstrates acceptable reliability with an alpha of 0.725.^32^ We included a seventeenth, intentionally false, statement, “Environmental factors, such as UV radiation, do not play a role in our genome.” This item was included to reflect the information in the skills module regarding environmental and genetic contributions to autism and was not included in calculating average knowledge scores.

#### Practical skills

Respondents read a one-page information sheet applying genetic information to human health, specifically genetic contributions to autism. This sheet included a cup-and-ball model presented in Hoang et al. (2018) that depicts how the combination of environmental and genetic factors can result in an ASD diagnosis. Respondents were then asked six questions about what they read and could refer to the information sheet as they responded to questions. The questions were designed based on those used in Abrams et al. (2015) and included both multiple-choice and fill-in-the-blank questions, such as “What is the purpose of genetic testing for autism spectrum disorder?” and “About what percentage of individuals who receive genetic testing are found to have a variant associated with higher risk for autism spectrum disorder?” Correctly answered questions received a score of 1 while all other question answers (e.g., incorrect or unsure responses) received a score of 0. Scores were calculated as a sum of correct answers, creating a final score range of 0 to 6. Based on Kuder-Richardson 20, the assessment is sufficiently reliable with an alpha of 0.654 based on 2021 GP data.^32^

#### Numeracy

Participants were asked to rate their ability to solve basic math problems on a scale from “1 - Not at all good” to “7 - Extremely good”, such as “How good are you at figuring out how much a shirt will cost if it is 25% off?” Scores were calculated as an average of all responses.^5^

#### ASD Diagnosis

Participants were asked if they have received an ASD diagnosis and responded with “Yes,” “No,” or “Not Sure.” This question was only included in the survey distributed to the SPARK sample.

#### ASD in Family

Participants were asked if someone in their family (e.g. parent, sibling, child) has been diagnosed with ASD, responding with “Yes,” “No,” or “Not Sure.”

#### Experience with ASD

Participants were asked broadly if someone in their life (e.g. friend, coworker, neighbor) has been diagnosed with ASD, responding with “Yes,” “No,” or “Not Sure.”

#### Education

Participants rated their highest level of education completed by selecting one of eight options, ranging from “No schooling completed” to “Doctorate degree.” These responses were used to create an ordinal variable dividing participants into four groups: “Less than high school,” “High school,” “Some college,” and “Bachelor’s or Graduate Degree.”

#### Age

Participants were asked to select their age from five groups: “18-25,” “26-39,” “40-49,” “50-59,” and “60+”.

#### Income

Participants were asked to report their income from three options: “Less than $49,999,” “$50,000-$99,999,” and “Over $100,000”.

### Statistical Analysis

Analyses were completed using R statistical software. To address our first aim, we conducted analyses of variance comparing scores in the three facets of genetic literacy (familiarity, knowledge, skills) between 2013 and 2021. Subsequent analyses controlled for education level given identified differences in self-reported education across the two samples. We used a propensity score adjusted comparison to control for education, in which we adjusted 2013 GP scores to match baseline variation in education in the 2021 GP. Adjusted mean scores are indicated by ‘2013 adj.’ and can be directly compared to mean scores in the 2021 GP.

To address our second aim, we conducted analyses of variance between scores in the three facets of genetic literacy assessed in the 2021 sample and the SPARK study sample. Given that there was no difference in education level between the 2021 GP and SPARK samples, we did not adjust for education in these analyses.

Exploratory analyses were conducted to determine potential associations between genetic literacy and other demographic and conceptual variables. We conducted simple correlations to determine associations, followed by a fitted regression model to determine significance.

To assess the replacement of the skills module within the genetic literacy measure, we replicated the analyses conducted in Abrams et al. (2015). This includes bivariate and partial correlations between the three facets of genetic literacy, controlling for education. We also fitted a regression model to determine if the skills segment mediates the relationship between familiarity and knowledge. Additionally, we conducted a Sobel test to determine the indirect effect between familiarity and knowledge, mediated by skills.

## Results

The characteristics of the 2021 GP sample (N = 2050) and SPARK sample (N = 2023) as compared to the 2013 GP sample are described in Table 1. The 2021 and SPARK samples self-report a higher average education level compared to the 2013 sample. The SPARK sample includes a higher proportion of individuals outside of the gender binary and a slightly younger sample on average.

**Table 1.** Sample Characteristics.

|  | 2013 GP | 2021 GP | SPARK |
| --- | --- | --- | --- |
| **Sample Size** | N = 1016 | N = 2050 | N=2023 |
|  | **%(N)** | **%(N)** | **%(N)** |
| **Gender** |  |  |  |
| Female | 52.3 (531) | 49.6 (1017) | 49.09(993) |
| Male | 47.7 (485) | 49.7 (1018) | 46.66(944) |
| Nonbinary | – | 0.56 (12) | – |
| Other | – | 0.15 (3) | 3.46(70) |
| Missing | – |  | 0.79(16) |
| **Race and Ethnicity** |  |  |  |
| Black or African American | 26.8 (272) | 27.5 (563) | 20.46 (414) |
| Latino or Hispanic American | – | 2.49 (51) | – |
| East Asian or Asian American | – | 4.58 (94) | 5.68(115) |
| South Asian or Indian American or Native Hawaiian | – | 1.07 (22) | 0.94(19) |
| Middle Eastern or Arab American | – | 0.25 (5) | – |
| White | 59.0 (599) | 57.0 (1169) | 51.90(1050) |
| Other | 2.4 (24) | 2.10 (43) | 5.34(108) |
| Multiple Selected | 3.7 (38) | 3.17 (65) | 14.88 (301) |
| Missing | – | – | 0.79(16) |
| **Age** |  |  |  |
| 18–25 | 15.8 (161) | 15.76 (323) | 8.06(163) |
| 26–39 | 23.7 (241) | 28.54 (585) | 43.3(876) |
| 40–49 | 36.2 (368) | 15.76(323) | 29.16(590) |
| 50–59 | 24.2 (26) | 11.8 (242) | 13.2(267) |
| 60+ | 15.8 (161) | 28.15 (577) | 5.49(111) |
| Missing | – | – | 0.79(16) |
| **Education Level** |  |  |  |
| Less than high school | 9.0 (91) | 5.17 (106) | 4.4(89) |
| High school | 23.2 (236) | 34.0 (697) | 26.5(536) |
| Some college | 29.3 (298) | 17.1 (350) | 20.91(423) |
| Bachelor’s or graduate degree | 30.2 (307) | 43.8 (897) | 47.4(959) |
| Missing | – | – | 0.79(16) |
| **Income** |  |  |  |
| Less than \$49,999 | 41.9 (426) | 49.5 (1015) | 45.8(926) |
| \$50,000–\$99,999 | 31.9 (324) | 31.1 (638) | 28.2(571) |
| Over \$100,000 | 26.2 (266) | 19.4 (397) | 25.2(510) |
| Missing | – | – | 0.79(16) |

Descriptive statistics of the three facets of genetic literacy within the three samples are presented in Table 2. We present the aggregate results within the three facets of genetic literacy and define “high literacy” as greater than 70% correct, as established in Abrams et al. (2015). As such, “high familiarity” indicates an average of at least 5 on the familiarity scale of 1 to 7. In the skills and knowledge segments, “high literacy” is defined as at least 5 out of 6 correct responses and at least 12 out of 16 correct responses, respectively. Average familiarity with the eight common genetic terms, as well as the ninth term “genome,” in each sample is presented in Figure 1. The 2021 sample on average reported moderate familiarity with the eight genetic terms. Participants were least familiar with the ninth term “genome” (M = 4.29, SD = 2.06) which was not included in the average familiarity score. The skills assessment cannot be directly compared to the 2013 data because different assessments and clinical examples were used. Participants in the 2021 GP sample correctly responded to approximately 3 of the 6 skills questions. Average scores for each of the six questions in the skills assessment are presented in Figure 2. The 2021 GP sample scored an average of 9 out of 16 on the genetic knowledge assessment compared to approximately 8 in 2013. Average scores for each question in the knowledge assessment, including the additional 17^th^ question, are presented in Figure 3.

**Figure 1.**
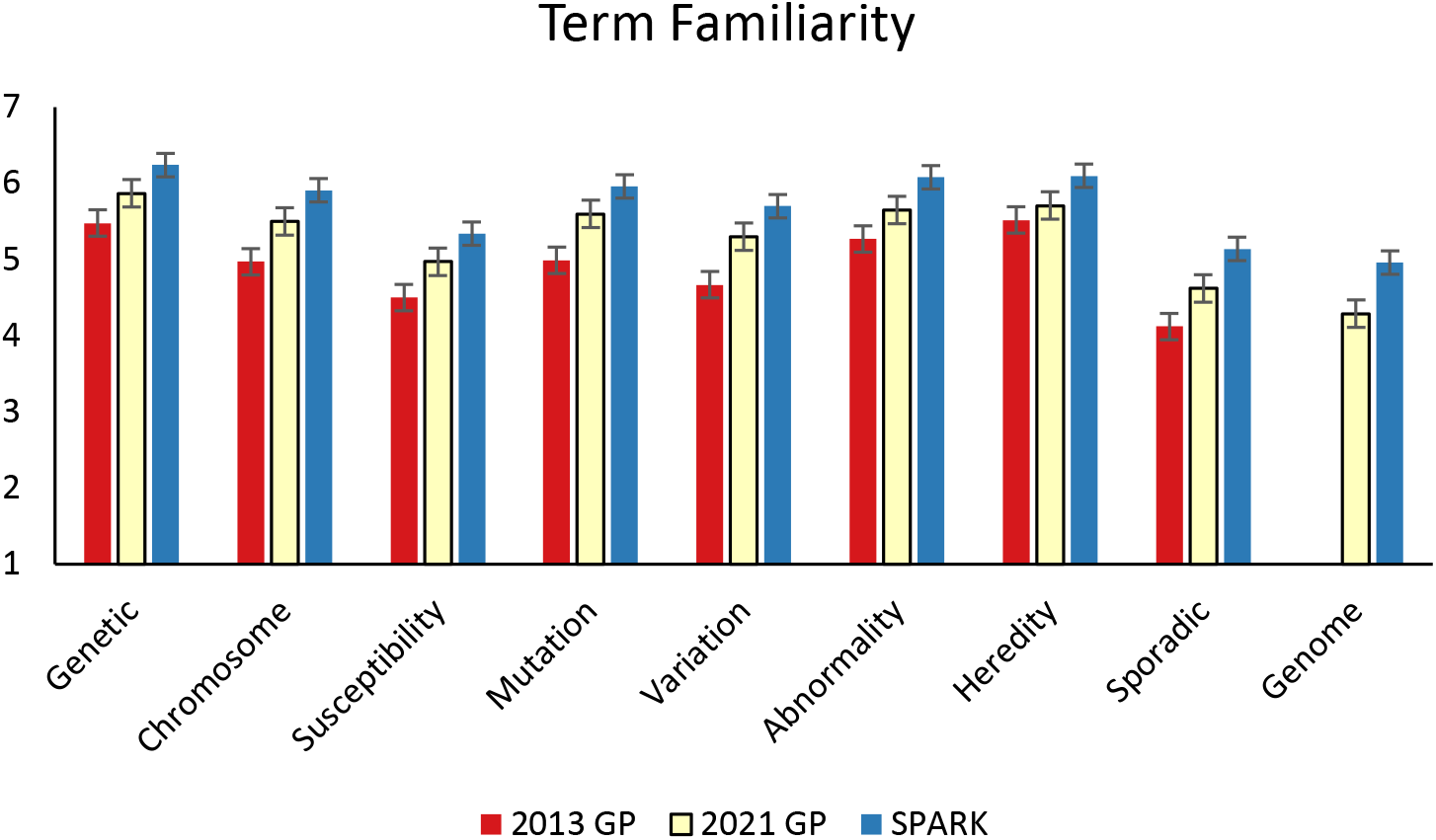
Participants’ self-reported familiarity with common genetic terms on a scale from 1 (not at all familiar) to 7 (completely familiar).

**Figure 2.**
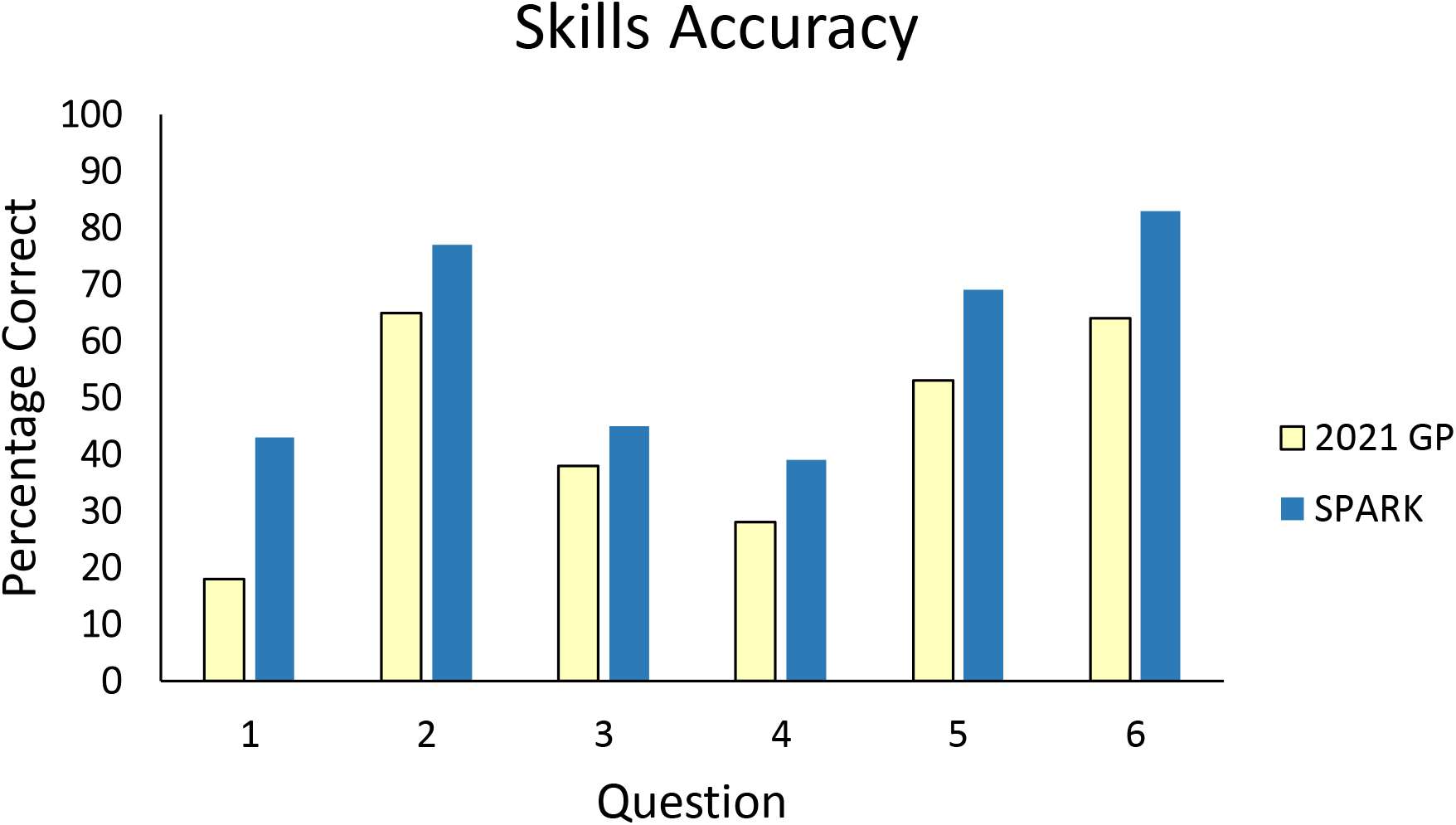
Percentage of each sample that correctly responded to each of six questions in the genetic literacy skills module.

**Figure 3.**
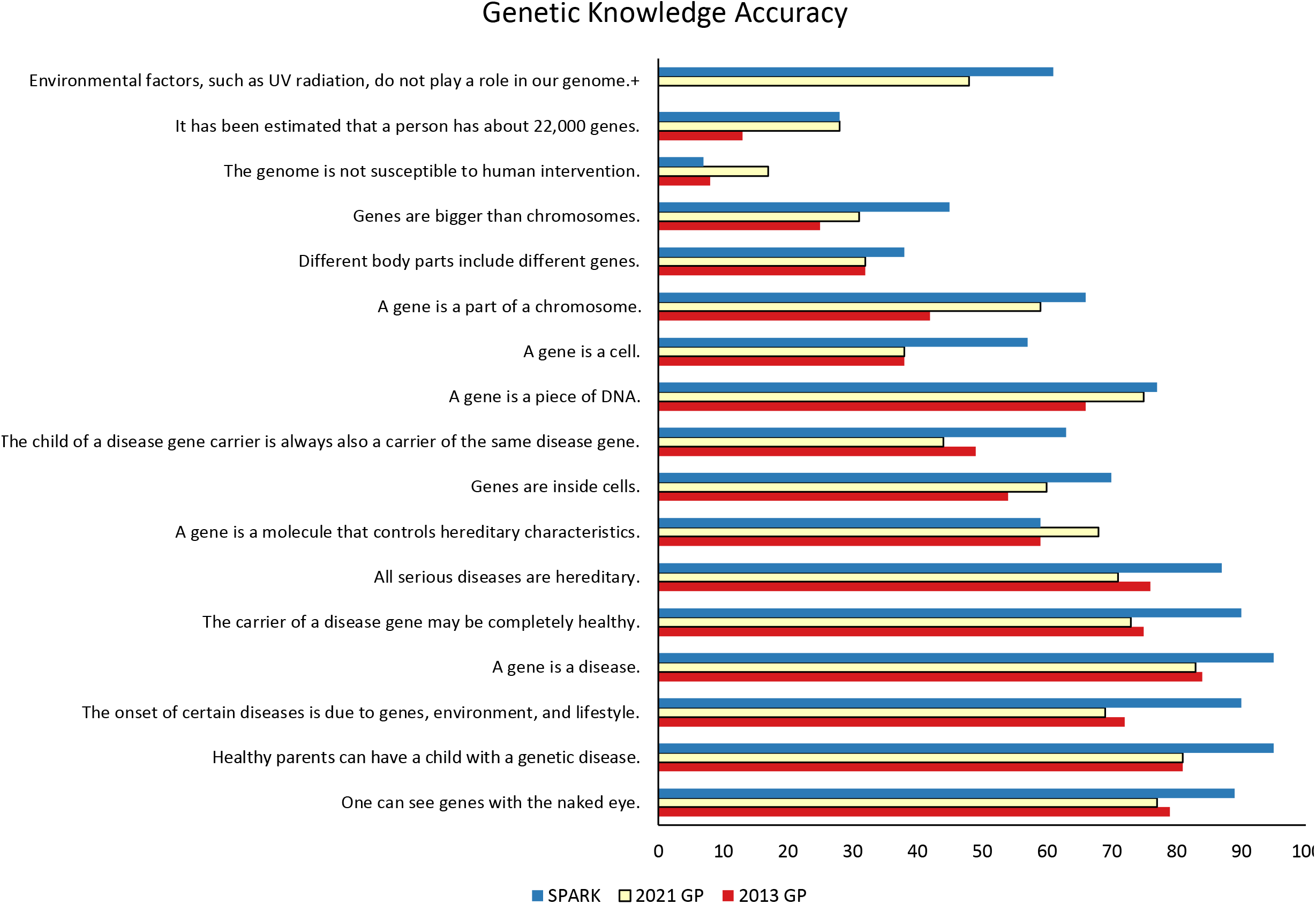
Percentage of each sample that correctly identifies each knowledge statement as true or false. +Not included in 2013 GP assessment.

**Table 2.** Descriptive statistics of the three genetic literacy measures in each sample.

|  | **Familiarity** | **Skills** | **Knowledge** |
| --- | --- | --- | --- |
| Range | 1–7 | 0–6 | 0–16 |
| **2013 General Population** |  |  |  |
| Mean(SD) | 4.98(1.76) | 3.94(1.88)* | 8.39(3.81) |
| % (N) Scoring>70% | 57.6 (585) | 47.5(483)* | 22.2(226) |
| **2021 General Population** |  |  |  |
| Mean(SD) | 5.36(1.36) | 2.65(1.73) | 9.06(3.23) |
| % (N) Scoring>70% | 66.9(1371) | 18.8(385) | 23.7(486) |
| **SPARK** |  |  |  |
| Mean(SD) | 5.79(1.27) | 3.57(1.56) | 10.57(2.96) |
| % (N) Scoring>70% | 79.4(1607) | 32.9(665) | 43.9(889) |

\*Different skills module implemented.

The SPARK sample also reported moderate familiarity, though higher than the 2021 GP sample. SPARK members correctly responded to 4 out of 6 skills questions, compared to 3 in 2021 (Table 2). Participants in the SPARK sample scored an average of 11 out of 16 points in the knowledge assessment (Table 2).

### General population updates in genetic literacy – 2013 to 2021

Average familiarity with genetic terms is significantly higher in 2021 (M = 5.40, SD = 1.39) compared to 2013 (M = 4.98, SD = 1.76, p <.001). After adjusting for variance due to education, the difference is still significant (2013 adj. M = 5.08, SD = 1.77, p<.001). Participants in the 2021 GP sample also reported significantly higher familiarity with each individual term compared to 2013 (Figure 1).

Scores in the knowledge assessment were significantly higher in the 2021 sample (M = 9.06, SD = 3.22) compared to 2013 (M = 8.52, SD = 3.82, p<.001). When adjusting for the variance due to education, the difference in average knowledge scores remains statistically significant (2013 adj. M = 8.72, SD = 3.76, p = 0.002). At the item level, the 2021 sample performed significantly better on seven of the sixteen items (p<.02, Figure 3). Some items showing significant improvement over the eight-year period include, for example, “It has been estimated that a person has about 22,000 genes,” (True) “A gene is a part of a chromosome,” (True) “A gene is a molecule that controls hereditary characteristics,” (True) and “The genome is not susceptible to human intervention” (False). The 2021 sample performed significantly lower on three of the sixteen items (p<.001, Figure 3). These statements are: “The onset of certain diseases is due to genes, environment, and lifestyle,” (True) “All serious diseases are hereditary,” (False) and “The child of a disease gene carrier is always a carrier of the same disease gene” (False).

### Differences in genetic literacy – general population versus research participants

Participants in the SPARK sample reported significantly higher familiarity with genetic terms overall (M = 5.79, SD = 1.27) compared to the 2021 general population (M = 5.40, SD = 1.39, p<.001). The SPARK sample also reported significantly higher familiarity with each of the eight genetic terms, as well as the ninth term “genome” (Figure 1). Knowledge scores were also significantly higher in the SPARK sample (M = 10.27, SD = 2.96) compared to the general population (M = 9.06, SD = 3.22, p<.001). A significantly higher proportion of the SPARK sample correctly determined the verity of 12 out of the 16 technical genetics statements (p<.001), compared to the 2021 general population (Figure 3). Example items include: “The onset of certain diseases is due to genes, environment, and lifestyle,” “All serious diseases are hereditary,” and “The child of a disease gene carrier is always also a carrier of the same disease gene.”

The SPARK sample also scored significantly higher on the skills module (M = 3.57, SD = 1.56) compared to the general population (M = 2.65, SD = 1.73, p<.001). Specifically, participants in the SPARK sample scored significantly higher in each of the six questions in the skills module (p<.001, Figure 2).

### Associations between genetic literacy and other variables – 2021 GP and SPARK

In the 2021 GP sample, participants’ self-reported numeracy significantly correlated with scores in familiarity (r = 0.34, p <.001), knowledge (r = 0.32, p<.001), and skills (r = 0.23, p<.001). In the SPARK sample, numeracy significantly correlated with familiarity (r = 0.33, p<.001) and knowledge (r = 0.38, p<.001) scores. In both samples, participants with a bachelor’s degree or higher scored significantly higher in all three facets compared to those who completed some or all of high school. Genetic literacy scores in both samples were higher in older participants and those with higher income.

Five hundred and ninety-three participants in the 2021 GP sample reported having an autistic person in their life, and these individuals scored higher in familiarity (M = 5.72, p<.001), knowledge (M = 9.77, p<.001), and skills (M = 2.96) than those without (M = 5.32; M = 8.97; M = 2.59).

In the SPARK sample, 838 individuals disclosed an autism diagnosis and scored marginally higher in all three facets of genetic literacy (M = 5.96; M = 10.88; M = 3.79) compared to non-autistic participants (M = 5.69; M = 10.43; M = 3.46). Those who reported knowing an autistic person in their life (N = 1323) also reported higher familiarity (M = 5.88) compared to those without (M = 5.65).

### Associations between genetic literacy measures and education

Results within each facet of genetic literacy in the 2021 sample positively correlated with each other and with education (Table 3). The same is shown for the SPARK sample. Partial correlations between the three facets of genetic literacy remained significant when adjusting for years of education.

**Table 3.**
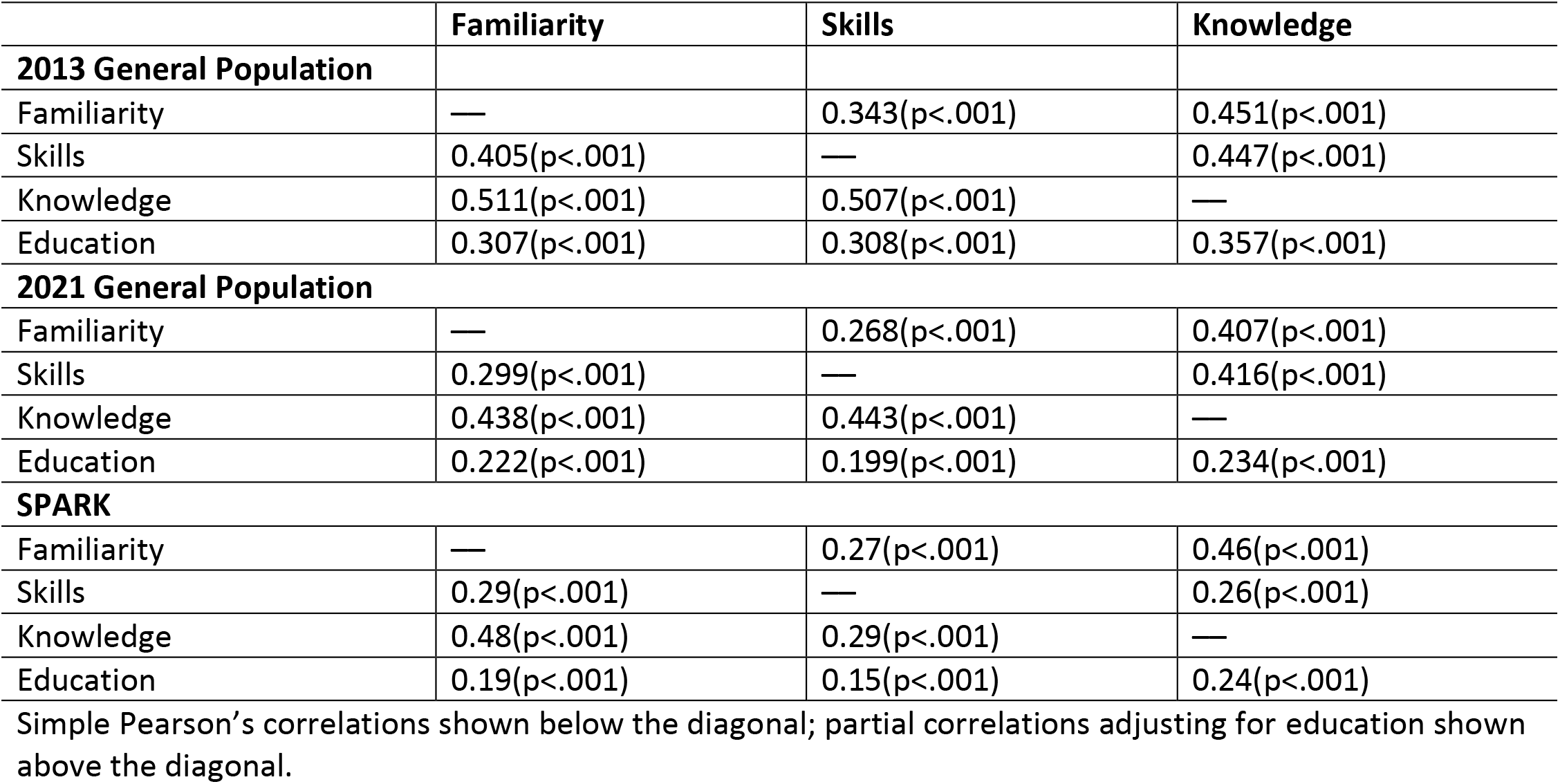
Pearson’s correlations of the three genetic literacy measures in each sample.

|  | **Familiarity** | **Skills** | **Knowledge** |
| --- | --- | --- | --- |
| **2013 General Population** |  |  |  |
| Familiarity | — | 0.343(p<.001) | 0.451(p<.001) |
| Skills | 0.405(p<.001) | — | 0.447(p<.001) |
| Knowledge | 0.511(p<.001) | 0.507(p<.001) | — |
| Education | 0.307(p<.001) | 0.308(p<.001) | 0.357(p<.001) |
| **2021 General Population** |  |  |  |
| Familiarity | — | 0.268(p<.001) | 0.407(p<.001) |
| Skills | 0.299(p<.001) | — | 0.416(p<.001) |
| Knowledge | 0.438(p<.001) | 0.443(p<.001) | — |
| Education | 0.222(p<.001) | 0.199(p<.001) | 0.234(p<.001) |
| **SPARK** |  |  |  |
| Familiarity | — | 0.27(p<.001) | 0.46(p<.001) |
| Skills | 0.29(p<.001) | — | 0.26(p<.001) |
| Knowledge | 0.48(p<.001) | 0.29(p<.001) | — |
| Education | 0.19(p<.001) | 0.15(p<.001) | 0.24(p<.001) |

Simple Pearson’s correlations shown below the diagonal; partial correlations adjusting for education shown above the diagonal.

We also fitted a regression model to test whether the skills segment mediates the relationship between familiarity and knowledge, while adjusting for education. Similar to the data in Abrams et al. (2015), the results were significant with a Sobel test^34^ indicating an indirect effect of familiarity on knowledge, mediated by skills (0.21, p<.001) (Figure 4).

**Figure 4.**
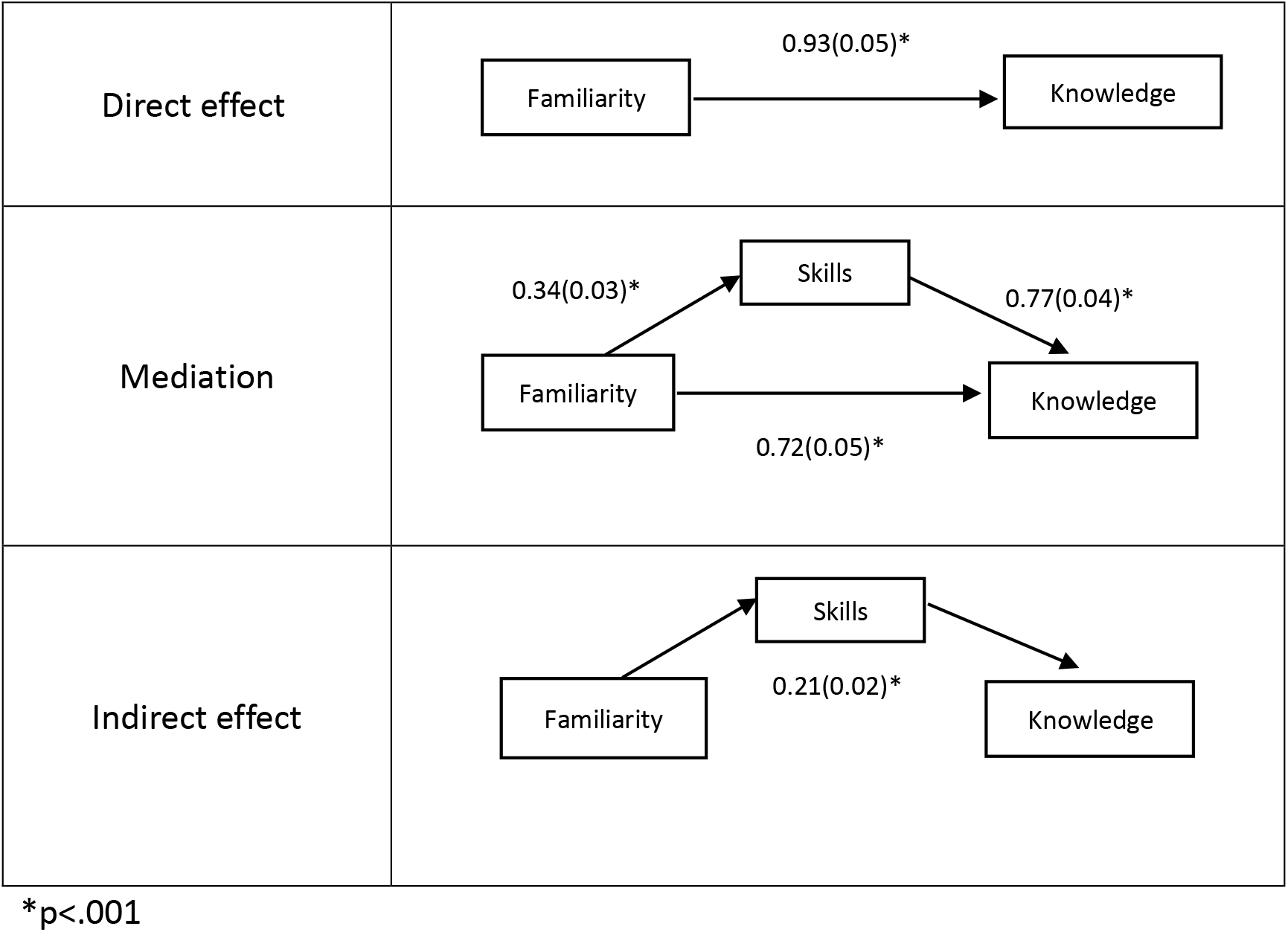
Meditation model

## Discussion

### Changes in Genetic Literacy Over Time

The survey data confirmed our first hypothesis that genetic literacy in 2021 is generally higher than 2013, reporting significantly higher knowledge of and familiarity with basic genetic concepts compared to Abrams et al. (2015). Considering the rapid advances in genomic research and utilization of genetic information in clinical and personal contexts, it follows that the general population’s genetic literacy will improve over time.^19^ In addition, depictions of genetics in the media, including celebrities undergoing genetic testing, the popularization of direct-to-consumer genetic testing and ancestry testing, and scientific fiction depicting genetic science only increase awareness of the impact of genetics on our lives.^5^ However, with the rise of genetic information, it is even more important to ensure that genetic education guides interpretation of genetic information in the media, to ensure misconceptions such as genetic determinism and genetic essentialism are not perpetuated.^35^ It is also necessary to equip the public with the skills and knowledge that will guide personal well-being and decision making. Though our data indicate higher genetic literacy in the current general population, it is important to explore significant changes or gaps in knowledge that have emerged in the past decade.

Participants in both the 2013 and 2021 general populations reported moderate familiarity with basic genetic terms, with the 2021 sample reporting significantly higher familiarity with each individual term and overall. These data support the notion that as the use of genetic testing and engagement with genomic research increases over time, so does general awareness of basic genetic terms. Like the 2013 sample, participants in 2021 were most familiar with general terms such as “genetic” and “heredity” and less familiar with risk terms such as “susceptibility” and “sporadic”. This suggests a need to expose the public to clinical applications of genetics and community-based genomics education programs in addition to basic technical terms.^36,37^ Participants in the 2021 sample were least familiar with the term “genome,” indicating a need for increased awareness of recent advances in genomic and personalized medicine. It is important to note, however, that familiarity with a term does not equate to comprehension and as such, it is important to also explore individuals’ factual knowledge.

On average, participants in the 2021 sample scored higher in the knowledge module, meaning they more accurately identified sixteen factual genetics statements as true or false. They were more accurate in identifying statements related to the definition and function of genes, such as “A gene is a molecule that controls hereditary characteristics” and “A gene is a piece of DNA,” indicating an improved understanding of basic biology. However, participants were also less accurate in identifying one statement as false, “The child of a disease gene carrier is always also a carrier of the same disease gene,” suggesting a need for skills in understanding inheritance patterns. This could be accomplished by applying knowledge of gene function to a clinical example, such as a family with genetic contributions to disease.

Somewhat surprisingly, the 2021 sample correctly responded to fewer than half of the skills questions, on average. This may be because of the high volume of text used in the infographic compared to that used in Abrams et al. (2015). Though both skills modules implemented six fill-in-the-blank or multiple-choice questions, the 2021 survey included one “select all that apply,” in which the correct response was to select more than one answer choice. We posit that this question added an element of difficulty not included in the Abrams et al. (2015) skills module and contributed to the reduction in skills scores. Given the different clinical context (i.e., hereditary breast and ovarian cancer vs. autism) and skills assessment across the two surveys, we cannot conclude from these data that participants’ genetic literacy skills have significantly changed between 2013 and 2021.

### Genetic Literacy in Genetic Research Participants

The results confirmed our second hypothesis that genetic literacy is higher in those enrolled in a large-scale genetic study. Compared to the general population sample recruited in 2021, those enrolled in SPARK reported significantly higher knowledge of and familiarity with basic genetic concepts, as well as an improved skills to synthesize information to make health decisions.

The SPARK sample reported moderate familiarity with basic genetic terms, though significantly higher than the 2013 and 2021 GPs for each term and overall. The SPARK sample also scored significantly higher in the knowledge segment on average. Moreover, the SPARK sample was significantly more accurate in determining the verity of more nuanced statements, such as “The onset of certain diseases is due to genes, environment, and lifestyle,” (True) and “The child of a disease gene carrier is always also a carrier of the same disease gene” (False).

This suggests that the SPARK sample is not only more aware of basic genetic concepts, but also has a better understanding of the interaction between genes and environments and the applications to health. Though the etiology of autism is not fully understood, current data show that it is caused by a complex interaction between genetics and environment, with previous studies indicating 70 to 90% heritability.^38,39^ Most notably, the SPARK sample performed significantly higher in the skills segment, both overall and within each question.

We posit that there are several reasons for the observed genetic literacy increase in the SPARK sample: first, participation in the SPARK study involves submission of medical information and genetic samples, along with access to multiple media resources on genetics generally and ASD specifically (examples at https://sparkforautism.org/discover/). Second, this sample’s experience with ASD, a highly heritable condition, may lead to improved recognition and comprehension of basic genetic concepts. This may be through interactions with the healthcare system, self-directed research, participation in advocacy groups, or all of the above. Similarly, a sample of 257 individuals with inflammatory bowel disease (IBD) in the US were asked to complete the same familiarity measure used here and as a group scored a 5.9 in familiarity out of 7 (completely familiar).^9^

Finally, beyond personal experience with a genetic condition, research experience alone is known to improve scientific literacy, and more specifically, genetic literacy. In the introductory biology classroom, data-based learning, or identifying biological concepts from previously implemented studies, promotes active learning and retention.^40^ Early introduction to research can also build students’ excitement for and success in science.^41^ Our data suggest that active participation in a research study, combined with personal experience with a genetic condition, can result in higher genetic literacy. Further research is needed to determine if this trend can be demonstrated in other genetic research participants.

Indeed, we chose to measure genetic literacy around autism in part because of the controversial nature of genetic research in the field.^42^ Members of the autism community have stated that they prioritize applied research (e.g., for increased daily supports) over basic research such as genetics^43^, and some autistics question whether the use of genetic research in ASD can be justified given past language dehumanizing autistics and justifying eugenic applications.^44^ We want to clearly state that any eugenic applications should be abhorrent to biomedical researchers and clinicians. We suggest that understanding of the facts and limitations of genetic research, along with true participatory engagement, is crucial to ensure any partnership with autistics and their families.

## Conclusion

Our results suggest that genetic literacy in the general population, specifically familiarity with genetic terms and demonstrable genetic knowledge, has improved in the past decade and that involvement in genetic research can improve one’s genetic literacy. However, there is still much room for improvement, particularly as genomic and personalized medicine becomes more mainstream. We recommend that educational interventions to improve genetic literacy are implemented in several domains: the general public, including school-aged children; patients affected by genetic or inherited conditions; and healthcare providers, highlighting both basic facts and the ability to apply information in multiple settings. In addition, educational interventions should focus on the ethical, legal, and social issues of contemporary genetics to combat misconceptions that can perpetuate harmful beliefs. The National Human Genome Research Institute also states improving genomic literacy as a significant priority, calling for active engagement between the public and genomic researchers.^45^ By equipping the public with the knowledge and understanding needed for personal health decisions, we are enabling advancements in genomic medicine to create positive change.

## Supplemental Information

Supplemental Information includes three tables and two spreadsheets.

## Acknowledgements

This work is funded by SPARK Research Match Diversity, Equity, and Inclusion (RM0149) and National Institutes of Health: Intramural Research Program of the National Human Genome Research Institute (HG200410-01). We would like to thank Daisy Yu of The Emmes Company, LLC for statistical assistance and Maddie Piper for insight into autistic perspectives.

## Data Availability

The published article includes the datasets collected as part of this study as supplementary files.

## Supplementary Tables

**Table S1.** Mean(SD) of familiarity with each genetic term on scale from 1 (not at all familiar) to 7 (completely familiar).

| Genetic Term | 2013 GP | 2021 GP | SPARK |
| --- | --- | --- | --- |
| Genetic | 5.48(1.83) | 5.87(1.47) | 6.26(1.2) |
| Chromosome | 4.97(2.01) | 5.50(1.66) | 5.94(1.41) |
| Susceptibility | 4.50(2.27) | 4.97(1.98) | 5.52(1.78) |
| Mutation | 4.99(2.07) | 5.59 (1.68) | 5.99(1.46) |
| Variation | 4.67(2.24) | 5.30(1.84) | 5.74(1.6) |
| Abnormality | 5.27(2.04) | 5.65(1.66) | 6.10(1.34) |
| Genome | — | 4.29 (2.06) | 5.00(1.87) |
| Heredity | 5.51(1.90) | 5.71(1.67) | 6.27(1.28) |
| Sporadic | 4.12(2.37) | 4.62(2.15) | 5.31(2) |

**Table S2.**
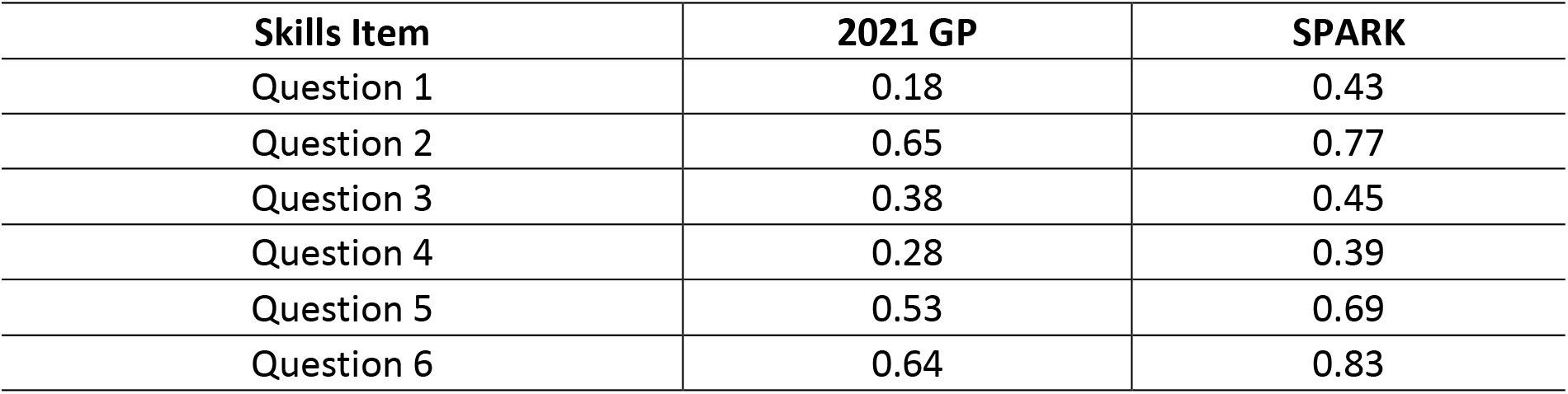
Proportion of the sample that correctly responded to each of six skills questions.

| Skills Item | 2021 GP | SPARK |
| --- | --- | --- |
| Question 1 | 0.18 | 0.43 |
| Question 2 | 0.65 | 0.77 |
| Question 3 | 0.38 | 0.45 |
| Question 4 | 0.28 | 0.39 |
| Question 5 | 0.53 | 0.69 |
| Question 6 | 0.64 | 0.83 |

**Table S3.** Proportion of each sample that correctly responded to each statement.

| Knowledge Statement | 2013 GP | 2021 GP | SPARK |
| --- | --- | --- | --- |
| **One can see genes with the naked eye.\*** | 0.79 | 0.77 | 0.89 |
| Healthy parents can have a child with a genetic disease. | 0.81 | 0.81 | 0.95 |
| The onset of certain diseases is due to genes, environment, and lifestyle. | 0.72 | 0.69 | 0.9 |
| **A gene is a disease.** | 0.84 | 0.83 | 0.95 |
| The carrier of a disease gene may be completely healthy. | 0.75 | 0.73 | 0.9 |
| **All serious diseases are hereditary.** | 0.76 | 0.71 | 0.87 |
| A gene is a molecule that controls hereditary characteristics. | 0.59 | 0.68 | 0.59 |
| Genes are inside cells. | 0.54 | 0.60 | 0.7 |
| **The child of a disease gene carrier is always also a carrier of the same disease gene.** | 0.49 | 0.44 | 0.63 |
| A gene is a piece of DNA. | 0.66 | 0.75 | 0.77 |
| **A gene is a cell.** | 0.38 | 0.38 | 0.57 |
| A gene is a part of a chromosome. | 0.42 | 0.59 | 0.66 |
| Different body parts include different genes. | 0.32 | 0.32 | 0.38 |
| **Genes are bigger than chromosomes.** | 0.25 | 0.31 | 0.45 |
| **The genome is not susceptible to human intervention.** | 0.08 | 0.17 | 0.07 |
| It has been estimated that a person has about 22,000 genes. | 0.13 | 0.28 | 0.28 |
| **Environmental factors, such as UV radiation, do not play a role in our genome.+** | — | 0.48 | 0.61 |

\***Bolded** statements intentionally false.
+Included only in survey completed by 2021 GP and SPARK.

## References

1. Knerr, S., Ramos, E., Nowinski, J., Dixon, K., and Bonham, V.L. (2010). Human difference in the genomic era: Facilitating a socially responsible dialogue. BMC Med Genomics 3, 20. 10.1186/1755-8794-3-20.

2. Incerti, D., Xu, X.M., Chou, J.W., Gonzaludo, N., Belmont, J.W., and Schroeder, B.E. (2022). Cost-effectiveness of genome sequencing for diagnosing patients with undiagnosed rare genetic diseases. Genet Med 24, 109–118. 10.1016/j.gim.2021.08.015.

3. Roberts, M.C., Allen, C.G., and Andersen, B.L. (2019). The FDA authorization of direct-to-consumer genetic testing for three BRCA1/2 pathogenic variants: a twitter analysis of the public’s response. JAMIA Open 2, 411–415. 10.1093/jamiaopen/ooz037.

4. Sabatello, M., Chen, Y., Sanderson, S.C., Chung, W.K., and Appelbaum, P.S. (2019). Increasing genomic literacy among adolescents. Genet Med 21, 994–1000. 10.1038/s41436-018-0275-2.

5. Abrams, L.R., McBride, C.M., Hooker, G.W., Cappella, J.N., and Koehly, L.M. (2015). The many facets of genetic literacy: Assessing the scalability of multiple measures for broad use in survey research. PLoS ONE. 10.1371/journal.pone.0141532

6. Kaphingst, K.A., Blanchard, M., Milam, L., Pokharel, M., Elrick, A., and Goodman, M.S. (2016). Relationships Between Health Literacy and Genomics-Related Knowledge, Self-Efficacy, Perceived Importance, and Communication in a Medically Underserved Population. Journal of Health Communication 21, 58–68. 10.1080/10810730.2016.1144661.

7. Milo Rasouly, H., Cuneo, N., Marasa, M., DeMaria, N., Chatterjee, D., Thompson, J.J., Fasel, D.A., Wynn, J., Chung, W.K., Appelbaum, P., et al. (2021). GeneLiFT: A novel test to facilitate rapid screening of genetic literacy in a diverse population undergoing genetic testing. J Genet Couns 30,, 742–754. 10.1002/jgc4.1364.

8. Meiser, B., Woodward, P., Gleeson, M., Kentwell, M., Fan, H.M., Antill, Y., Butow, P.N., Boyle, F., Best, M., Taylor, N., et al. (2022). Pilot study of an online training program to increase genetic literacy and communication skills in oncology healthcare professionals discussing BRCA1/2 genetic testing with breast and ovarian cancer patients. Fam Cancer 21, 157–166. 10.1007/s10689-021-00261-1.

9. Hooker, G.W., Peay, H., Erby, L., Bayless, T., Biesecker, B.B., and Roter, D.L. (2014). Genetic literacy and patient perceptions of IBD testing utility and disease control: a randomized vignette study of genetic testing. Inflamm Bowel Dis 20, 901–908. 10.1097/MIB.0000000000000021.

10. Chapman, R., Likhanov, M., Selita, F., Zakharov, I., Smith-Woolley, E., and Kovas, Y. (2019). New literacy challenge for the twenty-first century: genetic knowledge is poor even among well educated. J Community Genet 10, 73–84. 10.1007/s12687-018-0363-7.

11. Nakamura, S., Narimatsu, H., Katayama, K., Sho, R., Yoshioka, T., Fukao, A., and Kayama, T. (2017). Effect of genomics-related literacy on non-communicable diseases. J Hum Genet 62, 839–846. 10.1038/jhg.2017.50.

12. Haspel, R.L., Genzen, J.R., Wagner, J., Fong, K., and Wilcox, R.L. (2021). Call for improvement in medical school training in genetics: results of a national survey. Genet Med 23, 1151–1157. 10.1038/s41436-021-01100-5.

13. Kampourakis, K. (2017). Chapter 27 - Public Understanding of Genetic Testing and Obstacles to Genetics Literacy. In G. P. Patrinos (Ed.), Molecular Diagnostics (Third Edition) (Third Edit, pp. 469–477). Academic Press. https://doi.org/10.1016/B978-0-12-802971-8.00027-4

14. Alotaibi, A.A., and Cordero, M.A.W. (2021). Assessing Medical Students’ Knowledge of Genetics: Basis for Improving Genetics Curriculum for Future Clinical Practice. Adv Med Educ Pract 12, 1521–1530. 10.2147/amep.S337756.

15. Swandayani, Y.M., Cayami, F.K., Winarni, T.I., and Utari, A. (2021). Familiarity and genetic literacy among medical students in Indonesia. BMC Med Educ 21, 524. 10.1186/s12909-021-02946-8.

16. Schaibley, V.M., Ramos, I.N., Woosley, R.L., Curry, S., Hays, S., and Ramos, K.S. (2022). Limited Genomics Training Among Physicians Remains a Barrier to Genomics-Based Implementation of Precision Medicine. Front Med (Lausanne) 9, 757212. 10.3389/fmed.2022.757212.

17. Krakow, M., Ratcliff, C.L., Hesse, B.W., and Greenberg-Worisek, A.J. (2017). Assessing Genetic Literacy Awareness and Knowledge Gaps in the US Population: Results from the Health Information National Trends Survey. Public Health Genomics 20, 343–348. 10.1159/000489117.

18. Middleton, A., Milne, R., Almarri, M.A., Anwer, S., Atutornu, J., Baranova, E.E., Bevan, P., Cerezo, M., Cong, Y., Critchley, C., et al. (2020). Global Public Perceptions of Genomic Data Sharing: What Shapes the Willingness to Donate DNA and Health Data? Am J Hum Genet 107, 743–752. 10.1016/j.ajhg.2020.08.023.

19. Green, E.D., Gunter, C., Biesecker, L.G. et al. Strategic vision for improving human health at The Forefront of Genomics. Nature 586, 683–692 (2020). https://doi.org/10.1038/s41586-020-2817-4

20. Ishiyama, I., Nagai, A., Muto, K., Tamakoshi, A., Kokado, M., Mimura, K., Tanzawa, T., and Yamagata, Z. (2008). Relationship between public attitudes toward genomic studies related to medicine and their level of genomic literacy in Japan. Am J Med Genet A 146a, 1696–1706. 10.1002/ajmg.a.32322.

21. Kaphingst, K.A., Facio, F.M., Cheng, M.R., Brooks, S., Eidem, H., Linn, A., Biesecker, B.B., and Biesecker, L.G. (2012). Effects of informed consent for individual genome sequencing on relevant knowledge. Clin Genet 82, 408–415. 10.1111/j.1399-0004.2012.01909.x.

22. Erby, L.H., Roter, D., Larson, S., and Cho, J. (2008). The rapid estimate of adult literacy in genetics (REAL-G): a means to assess literacy deficits in the context of genetics. Am J Med Genet A 146a, 174–181. 10.1002/ajmg.a.32068.

23. Bowling, B.V., Acra, E.E., Wang, L., Myers, M.F., Dean, G.E., Markle, G.C., Moskalik, C.L., and Huether, C.A. (2008). Development and evaluation of a genetics literacy assessment instrument for undergraduates. Genetics 178, 15–22. 10.1534/genetics.107.079533.

24. Langer, M.M., Roche, M.I., Brewer, N.T., Berg, J.S., Khan, C.M., Leos, C., Moore, E., Brown, M., and Rini, C. (2017). Development and Validation of a Genomic Knowledge Scale to Advance Informed Decision Making Research in Genomic Sequencing. MDM Policy Pract 2. 10.1177/2381468317692582.

25. Linderman, M.D., Suckiel, S.A., Thompson, N., Weiss, D.J., Roberts, J.S., and Green, R.C. (2021). Development and Validation of a Comprehensive Genomics Knowledge Scale. Public Health Genomics 24, 291–303. 10.1159/000515006.

26. Feliciano P, Daniels AM, Snyder LG, et al. SPARK: a US cohort of 50,000 families to accelerate autism research. Neuron. 2018;97:488–93. 10.1016/j.neuron.2018.01.015

27. Kenny, L., Hattersley, C., Molins, B., Buckley, C., Povey, C., and Pellicano, E. (2016). Which terms should be used to describe autism? Perspectives from the UK autism community. Autism 20, 442–462. 10.1177/1362361315588200.

28. Bottema-Beutel, S.K.K., Jessica Nina Lester, Noah J. Sasson, and Brittany N. Hand (2021). Avoiding Ableist Language: Suggestions for Autism Researchers. Autism in Adulthood 3, 18–29. 10.1089/aut.2020.0014.

29. Gates, A.J., Gysi, D.M., Kellis, M., and Barabási, A.L. (2021). A wealth of discovery built on the Human Genome Project - by the numbers. Nature 590, 212–215. 10.1038/d41586-021-00314-6.

30. Jallinoja, P., and Aro, A.R. (1999). Knowledge about genes and heredity among Finns. New Genetics and Society 18, 101–110. 10.1080/14636779908656892.

31. Haga, S.B., Barry, W.T., Mills, R., Ginsburg, G.S., Svetkey, L., Sullivan, J., and Willard, H.F. (2013). Public knowledge of and attitudes toward genetics and genetic testing. Genet Test Mol Biomarkers 17, 327–335. 10.1089/gtmb.2012.0350.

32. Kuder, G. F., & Richardson, M. W. (1937). The Theory of Estimation of Test Reliability. Psychmetrika, 2, 151–160. https://doi.org/10.1007/BF02288391

33. Hoang, N., Cytrynbaum, C., & Scherer, S. W. (2018). Communicating complex genomic information: A counselling approach derived from research experience with Autism Spectrum Disorder. Patient Education and Counseling, 101(2), 352–361. https://doi.org/10.1016/j.pec.2017.07.029

34. Sobel, M.E. (1982) Asymptotic Confidence Intervals for Indirect Effects in Structural Equation Models. Sociological Methodology, 13, 290–321. https://doi.org/10.2307/270723

35. Donovan, B.M., Weindling, M., Salazar, B., Duncan, A., Stuhlsatz, M., and Keck, P. (2020). Genomics literacy matters: Supporting the development of genomics literacy through genetics education could reduce the prevalence of genetic essentialism. J. Res. Sci. Teach. 58, 520–550. https://doi.org/10.1002/tea.21670

36. Koehly, L.M., Morris, B.A., Skapinsky, K., Goergen, A., and Ludden, A. (2015). Evaluation of the Families SHARE workbook: an educational tool outlining disease risk and healthy guidelines to reduce risk of heart disease, diabetes, breast cancer and colorectal cancer. BMC Public Health 15, 1120. 10.1186/s12889-015-2483-x.

37. de la Haye, K., Whitted, C., and Koehly, L.M. (2021). Formative Evaluation of the Families SHARE Disease Risk Tool among Low-Income African Americans. Public Health Genomics 24, 280–290. 10.1159/000517309.

38. Johannessen, Jarle; Nærland, Terje; Bloss, Cinnamon; Rietschel, Marcella; Strohmaier, Jana; Gjevik, Elen; Heiberg, Arvid; Djurovic, Srdjan; Andreassen, Ole A.. Parents’ attitudes toward genetic research in autism spectrum disorder. Psychiatric Genetics: April 2016 - Volume 26 - Issue 2 - p 74–80. doi: 10.1097/YPG.0000000000000121

39. Genovese, A., and Butler, M. G. (2020). Clinical assessment, genetics, and treatment approaches in autism spectrum disorder (ASD). Int. J. Mol. Sci. 21, 1–18. doi: 10.3390/ijms21134726

40. Finby B, Heyer LJ, Malcolm Campbell A. Data-rich textbook figures promote core competencies: Comparison of two textbooks. Biochem Mol Biol Educ. 2021 May;49(3):392–406. doi: 10.1002/bmb.21488. Epub 2021 Jan 9. PMID: 33421340; PMCID: PMC8248048.

41. Jordan, T.C., Burnett, S.H., Carson, S., Caruso, S.M., Clase, K., DeJong, R.J., Dennehy, J.J., Denver, D.R., Dunbar, D., Elgin, S.C., et al. (2014). A broadly implementable research course in phage discovery and genomics for first-year undergraduate students. mBio 5, e01051–01013. 10.1128/mBio.01051-13.

42. Sanderson, K. (2021). High-profile autism genetics project paused amid backlash. Nature 598, 17–18. 10.1038/d41586-021-02602-7.

43. Roche L, Adams D, Clark M. Research priorities of the autism community: A systematic review of key stakeholder perspectives. Autism. 2021 Feb;25(2):336–348. doi: 10.1177/1362361320967790. Epub 2020 Nov 3. PMID: 33143455.

44. Botha, M., Dibb, B., and Frost, D.M. (2022). “Autism is me”: an investigation of how autistic individuals make sense of autism and stigma. Disability & Society 37, 427–453. 10.1080/09687599.2020.1822782.

45. Hurle, B., Citrin, T., Jenkins, J.F., Kaphingst, K.A., Lamb, N., Roseman, J.E., and Bonham, V.L. (2013). What does it mean to be genomically literate?: National Human Genome Research Institute Meeting Report. Genet Med 15, 658–663. 10.1038/gim.2013.14.

